# Acute muscle fatigue and CPR quality assisted by visual feedback devices: a randomizedcrossover simulation trial

**DOI:** 10.1101/399949

**Authors:** Cristian Abelairas-Gómez, Ezequiel Rey, Violeta González-Salvado, Marcos Mecías-Calvo, Emilio Rodríguez-Ruiz, Antonio Rodríguez-Núñez

## Abstract

**Objective:** To analyse the acute muscular fatigue (AMF) in triceps brachii and rectus abdominis during compression-only and standard cardiopulmonary resuscitation (CPR) performed by and certified basic life support providers.

**Methods:** Twenty-six subjects were initially recruited and randomly allocated to two study groups according to the muscles analysed; eighteen finally met the inclusion criteria (nine in each group). Both groups carried out two CPR tests (compression-only and standard CPR) of 10 min divided into five 2-min intermittent periods. The ventilation method was freely chosen by each participant (mouth-to-mouth, pocket-mask or bag-valve-mask). CPR feedback was provided all the time. AMF was measured by tensiomyography at baseline and after each 2-min period of the CPR test, in triceps brachii or rectus abdominis according to the study group.

**Results:** Rectus abdominis’ contraction time increased significantly during the fifth CPR period (p = 0.020). Triceps brachii’s radial muscle belly displacement (p = 0.047) and contraction velocity (p = 0.018) were lower during compression-only CPR than during standard CPR. Participants who had trained previously with feedback devices achieved better CPR quality results in both protocols. Half of participants chose bag-valve-mask to perform ventilations but attained lower significant ventilation quality than the other subjects.

**Conclusions:** Compression-only CPR induces higher AMF than standard CPR. Significantly higher fatigue levels were found during the fifth CPR test period, regardless of the method. Adequate rescuer’s strength seems to be a requisite to take advantage of CPR quality feedback devices. Training should put more emphasis on the quality of ventilation during CPR.

## Introduction

Cardiopulmonary resuscitation (CPR) is a physical activity that provokes fatigue in the rescuer. International guidelines for resuscitation promote two resuscitation protocols according to the scenario and the rescuers’ previous training: standard protocol [30 compressions & 2 rescue breaths] or compression-only CPR (continuous compressions) [1]. Physical fatigue caused by CPR in both protocols has been extensively documented [2,3]. Compression-only CPR (CO-CPR) produces more physical fatigue than standard CPR (Stand-CPR) [4,5]. However, fatigue has been generally estimated in terms of CPR quality [3-5]. In order to improve it, the use of feedback devices for learning and performing is increasing, although they may not necessarily reduce the effect of physical fatigue.

Muscle activation during CPR has been studied by electromyography [6,7,8]. However, we were unable to find any study that had examined acute muscular fatigue [AMF] as a consequence of different CPR protocols. To check neuromuscular fatigue, tensiomyography (TMG) has been identified as an objective, valid and reliable tool that allows evaluating the muscle contractile properties [9]. With this device, an electrical twitch stimulus is applied percutaneously and the consequent displacement caused by the muscle contraction is measured by a digital transducer pressed perpendicularly above the muscle belly. The displacement associated with the TMG response provides specific information on muscle tone or stiffness. Additionally, TMG measurements can be carried out quickly, without producing additional fatigue and do not depend on voluntary motivation [10].

Several studies have highlighted the usefulness and sensitivity of different TMG variables in detecting AMF following various kinds of exercise, such as ultra-endurance triathlon [9], strength training protocols [11], and eccentric exercise [12]. Generally, a loss of contractile properties has been observed by means of increased muscle contraction time and muscle tone, as well as decreased muscle contraction velocity [9,11,12]. Taking into the account the aforementioned arguments, TMG indices are expected to be able to illustrate the effects of CPR-related fatigue on mechanical capacities. This measure, together with other resuscitation variables, could provide a comprehensive picture of the effect of CO-CPR and Stand-CPR on acute fatigue.

The aim of our study was to analyse the AMF induced by good quality CPR performed by certified basic life support (BLS) providers.

## Materials and Methods

### Ethics

This randomized-crossover trial was carried out according to the Declaration of Helsinki. The study design was approved by the European University of the Atlantic Ethics Committee (Santander, Spain).

### Participants

Twenty-six people with basic life support (BLS) current certification (<6 months before starting the data collection) formed the initial study sample (convenience sample). They were asked to voluntarily participate in the study after being provided with details of its goals and methods. All of them were aged >18 and signed an informed consent form which further explained the aims of the research, the study design and the confidentiality statement. They were also informed that their participation was voluntary and they could withdraw at any time.

### Study design

A randomized-crossover trial was conducted from June to September 2017 (Figure 1). Body weight and height were measured with minimal clothing and bare feet. Age, handedness and usual performance physical exercise or not and type of training were recorded by oral request. Participants were randomly assigned to two groups using a random number generator: neuromuscular response was measured by TMG in triceps brachii in one group and in rectus abdominis in the second group.

**Fig. 1.**
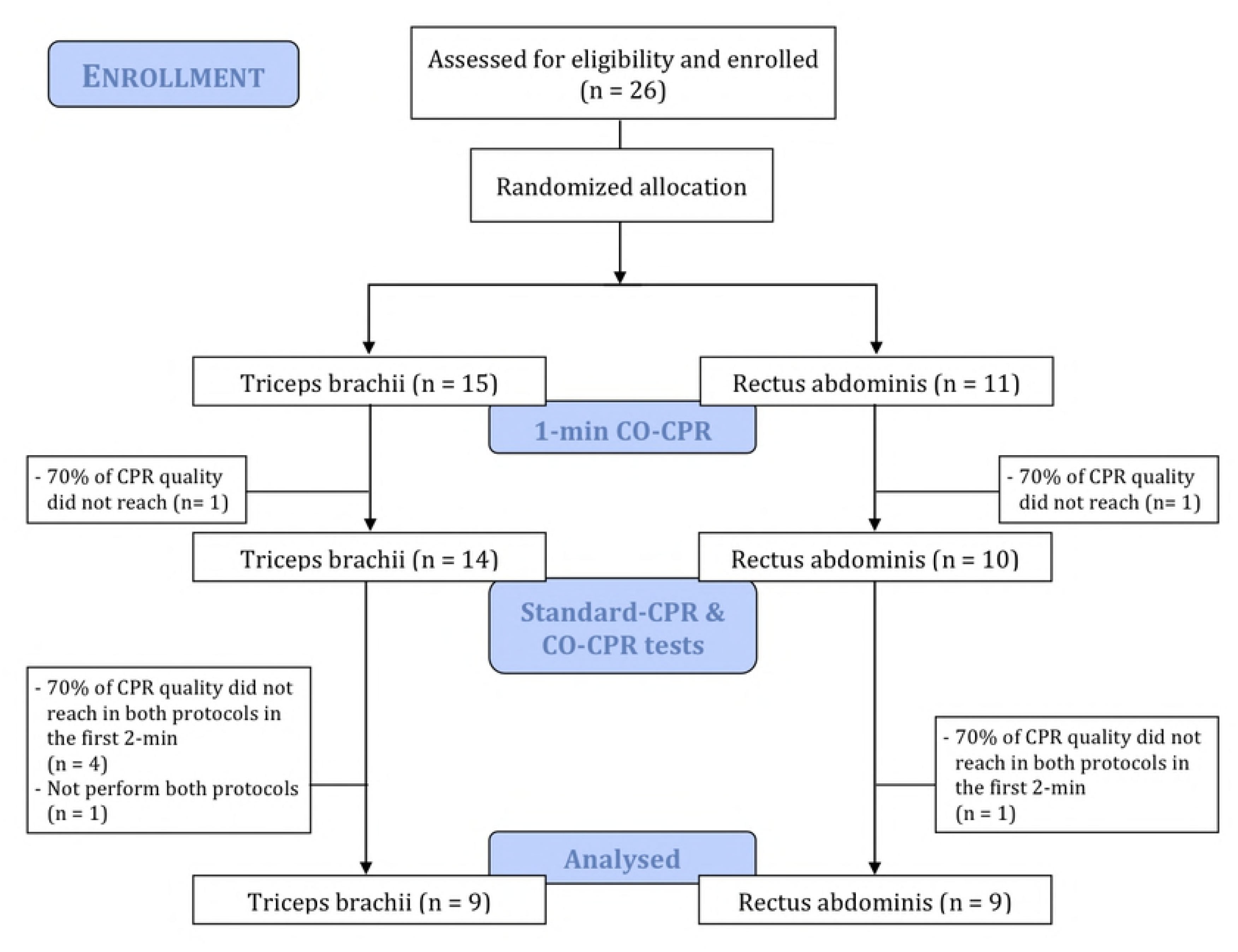
Study design. *Ventilation section* graphic with % of ventilations with excessive, insufficient and null volume and global ventilation quality (left); participants who achieved a mean of 500-600 ml (right). *Compression section:* analysis of both protocols, standard CPR (left) and compression-only CPR (right)

After randomization, participants performed an initial test consisting in one minute of CO-CPR on a sensitized manikin with visual feedback. Only if they were able to complete the test maintaining at least a 70% of quality in all the individually considered CPR variables (compression depth, compression rate, chest recoil and hands position), they were asked to continue the study. Participants who were not able to maintain such quality in this initial test were excluded.

Participants who continued the study were required to perform two additional CPR tests: Stand-CPR and CO-CPR, respectively. Two days of rest were left in between these tests and the order of performance was also randomized by means of a computer-generated list of random numbers. In both tests, a total time of 10 min of simulated CPR was divided into five 2-min periods, with a 2-min resting inter-period. Real-time visual feedback delivered by the manikin was provided. Participants who were not able to reach ≥70% of proficiency in all chest compressions’ quality parameters during the first 2 min of both protocols were excluded from the trial.

Prior to each test, a baseline measure with TMG and subsequent measures in each 2-min-resting period were conducted, with a total number of six TMG assessments in each CPR protocol. Finally, participants were asked about the CPR protocol that had subjectively produced them more physical fatigue (*“Regarding physical fatigue, which CPR-test was harder for you?”).*

### Acute muscle fatigue analysis

AMF of triceps brachii and rectus abdominis was assessed by TMG (TMG-S1 model). Measurements were performed under static and relaxed conditions before the CPR test and in the 2-min resting periods.

Briefly, TMG is composed of two electrodes through which a stimulation pulse of 1 ms and 0-100 mA is delivered. A displacement-measuring sensor situated between the electrodes records the changes in the muscle belly.

The sensor location was anatomically determined and marked with a dermatological pen. The sensor was pressed perpendicularly above the muscle surface. Electrodes of 5 x 5 cm were placed symmetric to the sensor. Increasing amplitudes of stimulation were delivered (50, 75 and 100 mA) [13], with a resting period of 15 seconds between consecutive measures to minimize the effect of fatigue and potentiation [14].

Variables analysed were maximum radial displacement in mm (Dm), contraction time (Tc) in ms, and contraction velocity (Vc) in mm·ms^-1^. Dm evaluates the muscle stiffness or tone and Tc corresponds to the time between 10% to 90% of Dm. Vc was calculated as Dm / (Tc + Td), where Td is the delay time, which corresponds to the time between the electric stimulation to 10% of Dm.

### CPR test and quality analysis

All CPR tests were performed on a Resusci Anne Manikin with PC Skillreporter Software (Laerdal, Norway), which provided CPR performance data. The manikin was configured according the European Resuscitation Council Guidelines for Resuscitation 2015 [1]. Rescue breaths could be delivered in three different ways: mouth-to-mouth (MtM), using a pocket mask or with bag-valve-mask (BvM). The method was freely chosen by the participant. Rescue breaths performed with null tidal volume because of incomplete airway opening and/or incorrect use of pocket mask/bag-valve-mask were also documented. In order to study the acute muscle fatigue caused by good quality resuscitation, participants were provided with the feedback delivered by the manikin in all tests.

### Data analysis

Shapiro-Wilk test was carried out to assess the normality of the variables. TMG variables were normally distributed, but CPR variables did not follow a normal distribution.

Tensiomyography data are presented as mean and standard deviation. Two intra-group factors (CPR-protocol: Stand-CPR vs. CO-CPR / CPR-period: 1 vs. 2 vs. 3 vs. 4 vs. 5) were analysed by repeated measures ANOVA in both muscles (triceps brachii & rectus abdominis). Partial eta squared (η^2^P) was calculated to measure the effect size. Mauchly’s Test of Sphericity was used to test the assumption of sphericity. When sphericity was not assumed, Greenhouse-Geisser correction was chosen to adjust the degrees of freedom.

Resuscitation variables are presented as median (Me) and interquartile rank (IQR). CPR-protocol and CPR-periods intra-group factors were also analysed. Inter-group factors were feedback-training (yes vs. no), physical-training (yes vs. no), ventilation method (MtM vs. BvM). Friedman test was used to assess intra-group differences at the five 2-min CPR-periods, with post-hoc Wilcoxon Signed Rank test to discern at which exact point significant differences occurred (significance level of p < 0.005). Inter-group analyses were performed using the Mann-Whitney U test. Effect size for non-parametric variables is reported using *r* and is interpreted as: small when *r* ≥ .10, medium when *r* ≥ 0.30, and large when *r* ≥ 0.50.

Statistical analyses were performed with IBM SPSS Statistics v.21 for Macintosh (v.21, Chicago, IL, USA). A significance level of p < 0.05 was considered for all analysis.

## Results

### Participants’ characteristics

Twenty-six BLS-certified subjects (12 females) were asked to participate in this study. After randomization, 11 and 15 participants were allocated to triceps brachii group and rectus abdominis group, respectively. Eighteen participants, 9 in each group, were able to achieve ≥70% of quality in all chest compression variables at the initial test and during the first 2 minutes of both protocols, and thus formed the final sample (Figure 1).

One participant was excluded because she only performed one CPR protocol. One male and five females were excluded for not meeting the abovementioned chest compression quality criteria; all exclusions were motivated by inadequate chest compressions’ depth: the man compressed too deep, while the five women did not reach the minimum depth. Of the excluded participants, only one had previously trained CPR skills with feedback, and only another affirmed practising exercise regularly.

Table 1 shows the baseline characteristics of the analysed sample (5 females). Only one participant was left-handed.

**Table 1.**
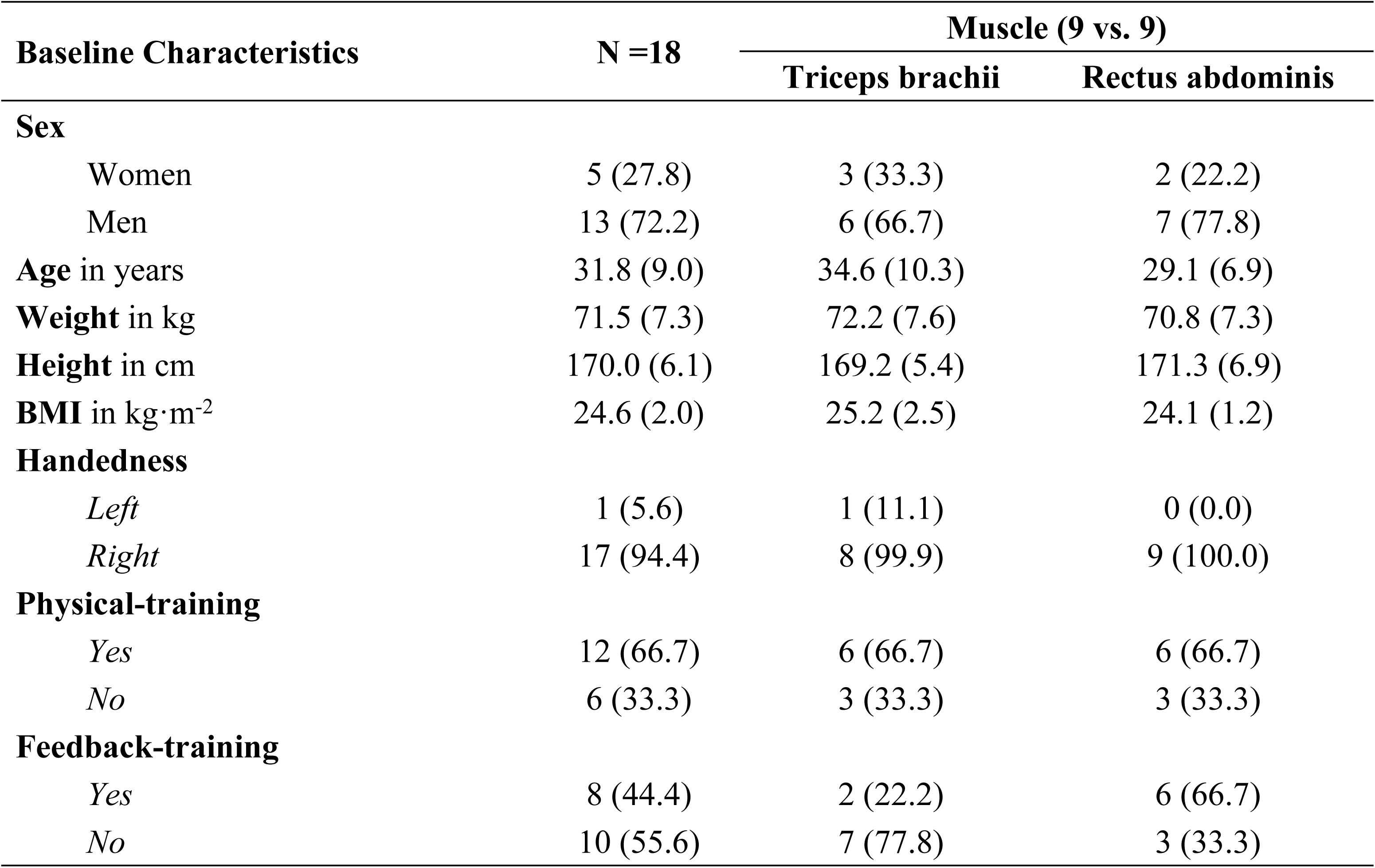
Participants’ characteristics. Categorical variables expressed as absolute frequencies (relative frequencies) and continuous variables as mean (standard deviation).

### TMG measurements

TMG data collection is presented in Table 2. No differences between CO-CPR and Stand-CPR were found in any variable (Tc, Dm and Vc) at baseline (p > 0.05 in all analysis).

**Table 2.** Analysis of TMG variables in the five 2 min CPR-periods. Results shown as mean (standard deviation). Analysis performed by repeated measures ANOVA.

| TMG Variables | | Period | | | | | Period-factor<br>p ($\eta^2_p$) |
| --- | --- | --- | --- | --- | --- | --- | --- |
|  |  | 1 | 2 | 3 | 4 | 5 |  |
| **Triceps brachii** |  |  |  |  |  |  |  |
| Tc | Stand-CPR | 15.8 (1.9) | 15.6 (2.4) | 15.7 (2.1) | 15.8 (2.0) | 16.2 (2.7) | 0.131 |
|  | HO-CPR | 15.9 (1.8) | 16.5 (4.2) | 15.9 (2.5) | 16.4 (2.3) | 16.6 (2.4) |  |
| | Test-factor | p ($\eta^2_p$) 0.153 | | | | | |
| Dm | Stand-CPR | 9.3 (2.7) | 8.0 (3.6) | 8.5 (2.8) | 8.1 (2.1) | 8.4 (2.3) | 0.321 |
|  | HO-CPR | 7.6 (2.6) | 6.8 (1.8) | 7.4 (1.7) | 7.0 (1.6) | 7.1 (2.1) |  |
| | Test-factor | p ($\eta^2_p$) 0.047 (0.409)* | | | | | |
| Vc | Stand-CPR | 0.27 (0.07) | 0.23 (0.08) | 0.25 (0.07) | 0.24 (0.05) | 0.25 (0.07) | 0.305 |
|  | HO-CPR | 0.22 (0.07) | 0.20 (0.05) | 0.21 (0.04) | 0.21 (0.04) | 0.20 (0.05) |  |
| | Test-factor | p ($\eta^2_p$) 0.018 (0.521)† | | | | | |
| **Rectus abdominis** |  |  |  |  |  |  |  |
| Tc | Stand-CPR | 37.5 (6.8) | 37.7 (6.8) | 38.9 (7.6) | 38.9 (7.1) | 41.2 (6.5) | A0.020‡<br>(0.567) |
|  | HO-CPR | 39.4 (5.4) | 38.3 (4.9) | 38.6 (4.8) | 38.2 (5.1) | 40.3 (4.7) |  |
| | Test-factor | p ($\eta^2_p$) 0.937 | | | | | |
| Dm | Stand-CPR | 10.7 (2.9) | 12.0 (2.8) | 11.6 (2.4) | 12.4 (3.0) | 12.2 (3.5) | 0.176 |
|  | HO-CPR | 11.0 (2.7) | 11.3 (1.5) | 11.2 (2.5) | 11.4 (2.0) | 11.6 (2.3) |  |
| | Test-factor | p ($\eta^2_p$) 0.628 | | | | | |
| Vc | Stand-CPR | 0.17 (0.06) | 0.20 (0.06) | 0.19 (0.05) | 0.20 (0.05) | 0.19 (0.06) | 0.257 |
|  | HO-CPR | 0.17 (0.04) | 0.18 (0.03) | 0.18 (0.04) | 0.18 (0.04) | 0.18 (0.04) |  |
| | Test-factor | p ($\eta^2_p$) 0.550 | | | | | |

\*: Significant differences found between test: Stand-CPR vs. HO-CPR, p = 0.047 ( $d = -0.622$ )
†: Significant differences found between test: Stand-CPR vs. HO-CPR, p = 0.018 ( $d = -0.790$ )
‡: Significant differences found between periods: Period 1 vs. Period 5, p = 0.008 ( $d = 0.470$ ); Period 3 vs. Period 5, p = 0.002 ( $d = 0.536$ ); Period 4 vs. Period 5, p = 0.043 ( $d = 0.411$ ).
^A: Sphericity not assumed. Adjusted by Greenhouse-Geisser.

Significant differences were found in the intra-group factor Period in Tc in rectus abdominis. In a further analysis, Tc in the fifth period was found to be higher than in the rest of periods (Table 2). Conversely, Tc did not vary significantly over time for triceps brachii. No differences were found when comparing CO-CPR and Stand-CPR in any muscle.

Dm and Vc were significantly lower in CO-CPR compared to Stand-CPR in triceps brachii. No differences were found in these variables in rectus abdominis.

### CPR performance

Regarding perceived physical fatigue, fourteen participants (77.8%) felt more fatigued after CO-CPR, three participants (16.7%) did not refer any difference and only one participant (5.6%) declared that Stand-CPR had been more exhausting.

### Quality of chest compressions

Positive and significant correlations were found between anthropometric variables and quality of chest compressions and mean depth. Height was positively correlated with global chest compressions’ quality (Stand-CPR: *r* = 0.697, p = 0.001; CO-CPR: *r* = 0.563, p = 0.015), mean depth (Stand-CPR: *r* = 0.485, p = 0.041; CO-CPR: *r* = 0.699, p = 0.001) and correct chest compressions’ fraction by depth (Stand-CPR: *r* = 0.595, p = 0.009; CO-CPR: *r* = 0.559, p = 0.016) in both protocols. Weight was positively correlated with mean depth (*r* = 0.525; p = 0.025) and correct chest compressions’ fraction by depth (*r* = 0.525; p = 0.025) in CO-CPR.

A descriptive analysis of compression variables is presented in Table 3. Quality of all chest compression variables was over 90% during the five 2-min periods. Mean depth was significantly different between Stand-CPR (Me: 54.0mm; IQR: 1.0) and CO-CPR (Me: 52.7 mm; IQR: 1.6; p = 0.005, *r* = 0.665), with no other differences between both protocols.

**Table 3.** Analysis of chest compression quality by 2-min CPR-period. Below, analysis of the feedback-training inter-group factor in the total 10-CPR. All results shown as median (interquartile rank).

| Variables | Period |  |  |  |  |
| --- | --- | --- | --- | --- | --- |
|  | 1 | 2 | 3 | 4 | 5 |
| Compressions quality |  |  |  |  |  |
| Stand-CPR | 95.6 (8.3) | 96.2 (6.6) | 95.5 (4.8) | 95.8 (6.1) | 95.4 (4.5) |
| HO-CPR* | 96.7 (5.2) | 97.0 (6.5) | 97.1 (5.9) | 95.9 (6.6) | 98.4 (4.4) |
| Mean rate |  |  |  |  |  |
| Stand-CPR† | 107 (10) | 111 (7) | 111 (6) | 114 (7) | 114 (6) |
| HO-CPR | 109 (8.5) | 114 (5.8) | 114 (6) | 115 (4) | 114 (5) |
| CCF by rate |  |  |  |  |  |
| Stand-CPR | 96.5 (8.3) | 99.0 (6.3) | 100.0 (4.5) | 98.0 (6.5) | 97.5 (7.0) |
| HO-CPR | 99.0 (4.5) | 99.0 (4.0) | 99.0 (3.5) | 99.0 (5.3) | 100.0 (1.3) |
| Mean depth |  |  |  |  |  |
| Stand-CPR | 54.0 (2.3) | 53.5 (3.3) | 53.5 (2.3) | 53.0 (2.3) | 53.5 (3.0) |
| HO-CPR | 52.5 (1.3) | 53.0 (1.3) | 53.0 (2.0) | 52.5 (3.0) | 53.0 (1.3) |
| CCF by depth |  |  |  |  |  |
| Stand-CPR | 93.5 (12.0) | 95.5 (9.5) | 94.0 (10.6) | 94.0 (14.5) | 94.0 (9.5) |
| HO-CPR | 96.5 (13.3) | 96.5 (9.3) | 97.5 (10.8) | 96.0 (11.3) | 99.0 (5.0) |
| CCF by chest recoil |  |  |  |  |  |
| Stand-CPR | 95.0 (10.5) | 96.5 (13.0) | 97.5 (8.0) | 98.0 (7.8) | 96.0 (7.3) |
| HO-CPR | 97.0 (7.0) | 97.0 (5.5) | 96.5 (6.3) | 97.5 (10.8) | 98.0 (5.3) |
| CCF by hands position |  |  |  |  |  |
| Stand-CPR‡ | 100.0 (2.3) | 100.0 (0.0) | 100.0 (0.0) | 100.0 (0.0) | 100.0 (0.0) |
| HO-CPR | 100.0 (0.0) | 100.0 (0.0) | 100.0 (0.3) | 100.0 (0.0) | 100.0 (0.0) |

**Chest compressions quality comparing feedback training with non-feedback training**
| Variables | Standard CPR |  |  | Compression-only CPR |  |  |
| --- | --- | --- | --- | --- | --- | --- |
|  | Feedback | N-Feedback | p (r) | Feedback | N-Feedback | p (r) |
| Compressions quality | 98.4 (2.8) | 93.1 (4.1) | 0.001<br>(0.754) | 99.1 (1.9) | 94.4 (6.9) | 0.003<br>(0.701) |
| Mean rate | 110 (8) | 111 (6) | 0.788<br>(0.063) | 113 (4) | 114 (7) | 0.398<br>(0.200) |
| CCF by rate | 98.7 (3.1) | 94.1 (6.4) | 0.021<br>(0.544) | 99.3 (1.9) | 98.2 (6.3) | 0.075<br>(0.418) |
| Mean depth | 54.0 (0.8) | 53.0 (2.0) | 0.049<br>(0.439) | 53.4 (1.3) | 52.6 (2.1) | 0.028<br>(0.513) |
| CCF by depth | 97.9 (5.8) | 86.4 (9.1) | 0.004<br>(0.680) | 98.3 (3.7) | 94.1 (17.6) | 0.021<br>(0.544) |
| CCF by chest recoil | 99.0 (7.2) | 92.8 (10.5) | 0.021<br>(0.544) | 98.6 (4.6) | 92.8 (6.6) | 0.045<br>(0.471) |
| CCF by hands position | 100.0 (0.0) | 100.0 (2.5) | 0.025<br>(0.418) | 100.0 (0.0) | 99.8 (1.5) | 0.011<br>(0.503) |

**Stand-CPR:** Standard cardiopulmonary resuscitation; **HO-CPR:** Hands-only cardiopulmonary resuscitation; **CCF:** Correct compressions fraction; **N-Feedback:** Non-Feedback.
Significance level between pairs of CPR-periods estimated in $p < 0.005$ .
\*: Significant differences found by Friedman test ( $p = 0.005$ ). Period 4 vs. Period 5, $p = 0.003$ .
†: Significant differences found by Friedman test ( $p < 0.001$ ). Period 1 vs. Period 4, $p = 0.002$ / Period 1 vs. Period 5, $p = 0.001$ .
‡: Significant differences found by Friedman test ( $p = 0.010$ ). No differences found between pairs of CPR-periods.

No differences were found in chest compression quality between participants who did exercise (n = 12) and sedentary participants (n = 6) in any protocol. In contrast, significant differences were found regarding previous feedback training (Figure 2). Participants who had previously trained CPR skills with feedback devices performed both protocols significantly better overall, except for mean rate (Stand-CPR & CO-CPR) and correct chest compressions’ fraction by rate (CO-CPR) (Table 3).

**Fig. 2.**
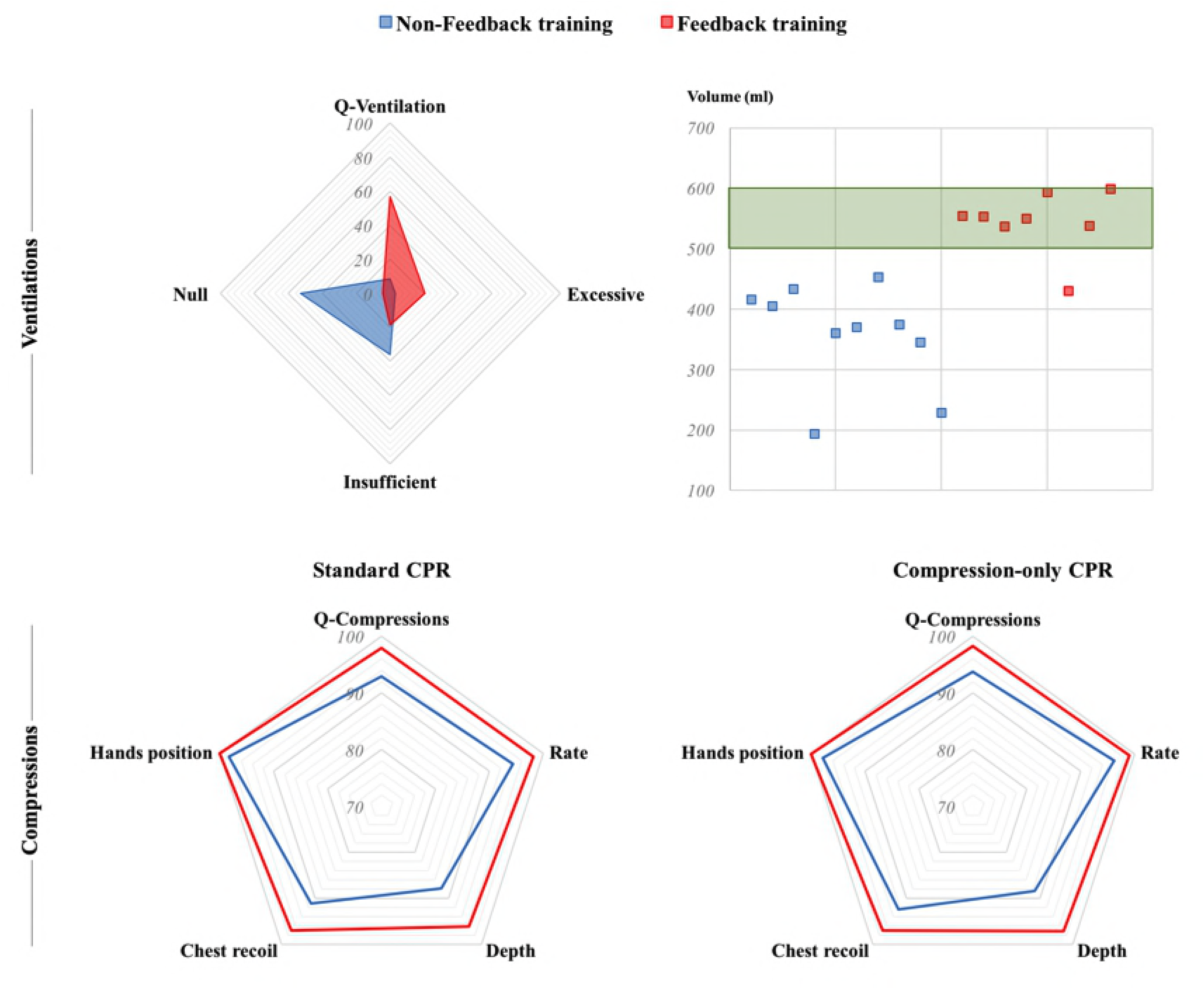
Comparison of CPR variables between participants with and without previous feedback training.

### Quality of rescue breaths

Nine participants selected MtM ventilation and 9 participants chose BvM. No differences were found in any variable in the intra-group factor CPR-period. Two factors were studied in the inter-group analysis: feedback-training and ventilation method (Table 4).

**Table 4.** Rescue breaths quality analysed from three inter-groups factors in the total 10-min CPR. Results shown as median (interquartile rank).

| Variables | Ventilation quality | Excessive volume | Correct volume | Insufficient volume | Null-tidal volume | No-flow Time | Tidal volume |
| --- | --- | --- | --- | --- | --- | --- | --- |
| **MtM**<br>(n = 9) | 50.0<br>(43.9-72.0) | 6<br>(2-17) | 25<br>(22-42) | 8<br>(5-14) | 0<br>(0-1) | 4.2<br>(4.0-5.6) | 550.6<br>(538-555) |
| **BvM**<br>(n = 9) | 3.9<br>(2.1-11.6) | 1<br>(0-2) | 2<br>(1-5) | 14<br>(13-20) | 27<br>(18-31) | 6.4<br>(6-7) | 370.2<br>(346-405) |
| *M-WU<sub>t</sub>* | 0.001 | 0.006 | 0.001 | 0.069 | 0.001 | 0.003 | 0.001 |
| *r* | 0.801 | 0.645 | 0.791 | 0.427 | 0.739 | 0.708 | 0.760 |
| **Feedback**<br>(n = 8) | 60.9<br>(44.0-75.6) | 7<br>(4-19) | 31<br>(24-42) | 8<br>(5-13) | 0<br>(0-1) | 4.2<br>(4.0-5.2) | 552.2<br>(538-575) |
| **Non-Feedback**<br>(n = 10) | 6.3<br>(2.1-12.5) | 1<br>(0-2) | 3<br>(1-6) | 16<br>(13-20) | 26<br>(18-31) | 6.4<br>(6-7) | 372.8<br>(346-416) |
| *M-WU<sub>t</sub>* | 0.001 | 0.005 | 0.001 | 0.032 | 0.001 | 0.003 | 0.001 |
| *r* | 0.796 | 0.660 | 0.792 | 0.503 | 0.796 | 0.691 | 0.796 |

**MtM:** Mouth-to-mouth; **BvM:** Bag-valve-mask; **M-WU<sub>t</sub>:** Mann–Whitney U test.
Ventilation quality in %; Excessive, correct, insufficient and null-tidal volume shown as frequencies; No-flow time in seconds; Tidal volume in ml.

All participants who had previously received feedback-training selected MtM (n = 8) and achieved better results in all variables. Percentage of correct rescue breaths was statistically higher without barrier device (MtM, Me: 52.1%, IQR: 46.4; BvM, Me: 3.9%, IQR: 10.0; p = 0.001, *r* = 0.801). The use of BvM implied a greater number of null-tidal volume ventilations, more no-flow time and less tidal-volume.

Figure 2 shows differences in rescue breaths’ quality between participants who had previously trained with feedback and those who had not. Participants with no previous feedback training were unable to reach a mean tidal volume of 500-600 ml in the total 10-min CPR test.

## Discussion

During CPR practise, triceps brachii plays an important role in extending the elbow [15], whereas rectus abdominis controls the trunk and stabilizes the upper body to provide a proper force distribution during chest compressions [6]. For this reason, contractile properties of these muscles were measured through non-invasive and non-demanding TMG in this study.

While the selected CPR protocols did not entail severe symptoms of fatigue during the four first 2 min periods, during the last set [period 5] acute fatigue symptoms were observed in both protocols, with a sudden significant increase of Tc for rectus abdominis, causing greater latency for muscle contraction. Despite comparisons are difficult due to strongly contrasting forms of induced fatigue, the increase in muscle contraction time observed in our study is in agreement with previous investigations analysing the influence of different types of exercise on TMG muscle properties [9,11,16]. This may be partially explained by a reduced efficiency of the excitation-contraction coupling, impaired membrane conduction properties and destruction of cellular structures (i.e. peripheral fatigue) [11]. Based on the present findings, whatever CPR protocol is applied, a 2-min resting period appears to be insufficient to recover at least rectus abdominis mechanical properties after 4 consecutive 2-min periods of simulated CPR.

The present data also show that the Tc for rectus abdominis and triceps brachii was longer than 30ms, indicating they are highly resistant to fatigue and with a high prevalence of slow-twitch fibres [17]. This data may have important implications to design potential muscle-training programs for CPR professionals.

Differences in fatigue levels and changes in body biomechanics between CO-CPR and Stand-CPR have been previously reported [7]. As Trowbridge et al. reported significantly greater perceived effort and joint torque changes in CO-CPR [7], the differences in Dm and Vc between CO-CPR and Stand-CPR found in this study were consistent with our expectations. Using TMG, we found that after each CPR period the CO-CPR group showed significantly lower values in Dm (greater stiffness) and Vc (slower muscle fibre conduction) in triceps brachii and hence, greater signs of fatigue than Stand-CPR group. Despite the precise reasons for Dm and Vc reduction in the CO-CPR are unclear, there are a number of possible explanations, such as impairments in excitation-contraction coupling, loss of membrane conduction properties and cellular structures destruction, which in turn result in increased muscle tone and/or reduction in the muscle’s ability to generate force [18].

Acute muscular fatigue was higher during CO-CPR. Taking into account these results, it is plausible that 2-min cycles of continuous chest compressions could induce too much neuromuscular fatigue, compromising CPR quality. A previous study concluded that fatigue of the spinal and lumbar musculature measured by electromyography occurred after 2-min. In this case, participants had to reach an 80% of chest compression quality without feedback, and those who failed to do it were excluded [8]. In our study, AMF had no influence on performance quality, maybe because participants were guided by visual feedback, which helped to ensure adequate CPR quality and allowed studying the AMF this produced. However, further studies without real-time feedback should be conducted to determine the impact of acute fatigue on a real scenario.

CPR is a physically demanding activity that requires certain levels of coordination and strength. In our study, seven participants were excluded because they were not able to reach a good compression quality despite feedback. Additionally, six of them did not usually exercise, and it is known that some workouts such as strength training programs may be useful to reach and maintain chest compressions’ quality [19]. Considering the participants finally enrolled, no differences were found between those who did exercise and not, this might be due to the reduced number of subjects who completed the study.

As it has been formerly reported by other studies and according to our results, anthropometric measurements correlate with the capacity to compress deeper [2,20,21], thus greater body sizes could compensate low strength levels.

Although the use of feedback devices might not be sufficient to deliver high quality compressions in case of strength shortage, our results showed their effectiveness in a simulated situation when participants were in adequate physical condition. Thus, compression quality was over 90% in all periods in both protocols. This was not the case of rescue breaths, whose quality remained below 70%, especially for participants without previous feedback-enhanced training and for those who decided to use BvM (<10% in both cases). Low ventilation quality during CPR, as well as differences between MtM and BvM ventilation have already been described, but usually without use of feedback devices [22].

### Limitations

This is to our knowledge the first study attempting to analyse physical fatigue caused by CPR with TMG. Although TMG has been reported as a quick and non-invasive method to describe muscular properties, more research is needed to characterize and fully understand the amount of data that it is able to register.

CPR quality might have been overestimated in our analysis. In real conditions, feedback is usually not available to help the rescuer and adjust his/her performance. On the other hand, the use of feedback during tests might have underestimated the effect of AMF on CPR quality.

## Conclusions

AMF induced by compression-only CPR was higher than that induced by standard CPR, and this was also higher in the fifth 2-min CPR period comparing to the previous. However, participants were able to achieve good CPR quality despite AMF. More studies are needed to clarify if the AMF found or an inadequate technique could decrease CPR quality without feedback devices, which might not guarantee CPR quality if the rescuer’s physical strength is insufficient. Ventilation quality needs to be reinforced during CPR training, with special emphasis on the BvM procedure.

